# Identification of human genetic variants controlling circular RNA expression

**DOI:** 10.1101/472977

**Authors:** Ikhlak Ahmed, Thasni Karedath, Fatima M. Al-Dasim, Joel A. Malek

## Abstract

Circular RNAs (circRNAs) are abundant in eukaryotic transcriptomes and have been linked to various human disorders. However, understanding genetic control of circular RNA expression is in early stages. Here we present the first integrated analysis of circRNAs and genome sequence variation from lymphoblastoid cell lines of the 1000 genomes project. We identified thousands of circRNAs in the RNA-seq data and show their association with local single nucleotide polymorphic sites, referred to as circQTLs, which influence the circRNA transcript abundance. Strikingly, we found that circQTLs exist independently of eQTLs with most circQTLs having no effect on mRNA expression. Only a fraction of the polymorphic sites are shared and linked to both circRNA and mRNA expression with mostly similar effects on circular and linear RNA. A shared intronic QTL, rs55928920, of HMSD gene drives the circular and linear expression in opposite directions, potentially modulating circRNA levels at the expense of mRNA. Finally, circQTLs and eQTLs are largely independent and exist in separate linkage disequilibrium (LD) blocks with circQTLs highly enriched for functional genomic elements and regulatory regions. This study reveals a previously uncharacterized role of DNA sequence variation in human circular RNA regulation.

## Introduction

Circular RNAs (circRNAs) are an abundant class of regulatory transcripts primarily derived from protein coding exons and widely expressed across eukaryotic organisms including *Homo sapiens* and *Mus musculus* [1–6]. These transcripts are produced co-transcriptionally; the 5’ and 3’ are joined in a head-to-tail “backsplice” junction to form a circular molecule leaving no exposed ends. CircRNAs are known to be evolutionarily conserved across species, relatively stable in the cytoplasm and often show tissue/developmental stage-specific expression pattern. CircRNAs have been shown to be involved in post-transcriptional regulation by functioning as decoys for the binding of miRNAs, reducing their ability to target mRNAs [7,8]. Specifically, the circRNA ciRS-7, also known as CDR1as, has been found to harbor over 70 miR-7 binding sites and functions as an miRNA sponge to repress miR-7 activity. Cdr1as knockout mice suffer from defects in sensorimotor gating [9] and silencing of CDR1as in colorectal cancer and hepatocellular carcinoma cell lines resulted in decreased tumor proliferation [10,11]. Similarly, the circular RNA of the Sry gene contains 16 binding sites for miR-138, and miR-138 mediated knockdown of mRNA was shown to be attenuated with circular Sry overexpression [7]. Indeed, multiple studies have reported the sequestration of miRNAs by circRNAs, making them excellent candidates for competing endogenous RNA activity [7,8,12–16]. Although multiple reports have emerged indicating certain circRNAs function as miRNA sponges, increasing evidence indicates that circular RNAs may have other potential functional roles like storing or sequestering transcription factors or RNA binding proteins [17], microRNA transport [9] and at least some are translatable into functional proteins [18–21]. Legnini et al. [21] were able to show that circ-ZNF609 is translated into a protein in a splicing-dependent and cap-independent manner. The resultant protein has a functional role in muscle differentiation in Duchenne muscular dystrophy by regulating myoblast proliferation. circRNAs have been shown to be altered in a variety of pathological conditions, which has stimulated significant interest into their role in human disease and cancer [13,22]. Although their exact roles and mechanisms of gene regulation remain to be understood, circRNAs are being actively investigated for their association with diseases and their role as potential biomarkers and novel therapeutic targets.

Recent association studies on expression quantitative trait loci (eQTL) have provided information on genetic factors, especially single nucleotide polymorphisms (SNPs), associated with variation in gene expression. These studies have demonstrated the regulatory role of SNPs on gene expression and the splicing patterns of mRNA [23–26]. Expression QTL studies offer an excellent platform to link DNA sequence variation to changes in gene expression that may contribute to phenotypic diversity and disease susceptibility.

In this study we present the first integrated analysis of genome sequence variation and circular RNA expression to identify a set of regulatory variants influencing circRNA expression. We identified thousands of circRNAs in RNA-seq data of lymphoblastoid cell lines from the 1000 genomes project [24] and mapped their association with local single nucleotide polymorphic sites, referred to as circQTLs. These circQTLs associate with the circRNA abundance and exist independently of eQTLs with most circQTLs having no effect on mRNA expression. We report that, only a fraction of the polymorphic sites are shared and linked to both circRNA and mRNA expression with mostly similar effects on circular and linear RNA. We also show that circQTLs and eQTLs exist in separate linkage disequilibrium (LD) blocks with circQTLs enriched for exonic and regulatory region variants. This study reveals a previously uncharacterized role of DNA sequence variation in human circular RNA regulation.

## Results

### Assessing circRNA expression

We used RNA sequencing data of 358 lymphoblastoid cell line (LCL) samples of European ancestry sequenced in the framework of Geuvadis RNA-sequencing project [24] along with their genotype information from 1000 Genomes data [27]. We analyzed circRNA expression of these samples using our in-house developed circular RNA detection pipeline [28]. This pipeline identifies circular RNA structures by using Bowtie 2 [29] to align paired-end RNA-seq data to a custom reference exome containing all possible pairs of intragenic non-linear combinations of exons, as well as single exon “backsplices”. This process yielded a total of 95675 unique circRNA candidates with a minimum of 50 independent junctional reads across all samples and expression in at least 60% of the samples (Supplementary_File_S1.xlsx). Of the 95675 circRNAs as identified by the pipeline, 58050 overlap with one or more circRNA loci defined in the circBase [30] (Supplementary_File_S2.txt).

We have validated several circRNA candidates identified in this study in Human T lymphocyte Jurkat cells using multiple approaches. Divergent primers were designed for a set of circRNAs to perform PCR amplifications of the backsplice exon junction from cDNA of the Human T lymphocyte Jurkat cells (Supplementary_File_S3.xlsx). An “outwad-facing” strategy in the design of the divergent primers guarantees the amplifications are from a circular template [31]. For all the 17 circRNA candidates tested, a single distinct band of expected product size was obtained in a PCR assay and the Sanger sequencing of the PCR products confirmed the presence of the backsplice junction sequence (Figure 1, Supplementary_Figure_1.pdf). As a control, we also designed convergent primers to amplify the linear RNA (mRNA) of the respective host genes of some of the circRNA candidates. Because the mRNA structure is dictated by the genomic order of exons, PCR primers designed to amplify the linear RNA can produce bands for both cDNA and genomic DNA (gDNA) fractions. On the other hand, the backsplice junction sequence of the correctly identified circRANs should not exist in the genome and hence circRNA specific primers should only be able to amplify in the cDNA but not the gDNA fraction. Indeed, as expected, positive and negative results of amplifications were obtained for cDNA and gDNA fractions for circRNA specific primers. Primers designed to amplify linear RNA produced bands of expected product sizes for both cDNA and gDNA (Figure 1C). Further evidence of a circularized structure for these circRNA candidates came from the digestion of total RNA with an exoribonuclease enzyme RNase R. This exonuclease enzyme digests all linear RNA forms with a 3’ single stranded region of greater than 7 nucleotides [32]. The circRNAs have been shown to resist the RNase R mediated digestion due to their lack of 3’ single strand overhangs and hence show enrichment in the RNase R treated samples. Indeed, there was ample enrichment of our tested circRNA candidates after the RNase R treatment compared to mRNA, confirming their resistance to the exoribonuclease digestion and strongly attesting to their non-linear structure (Figure 1D).

**Figure 1.**
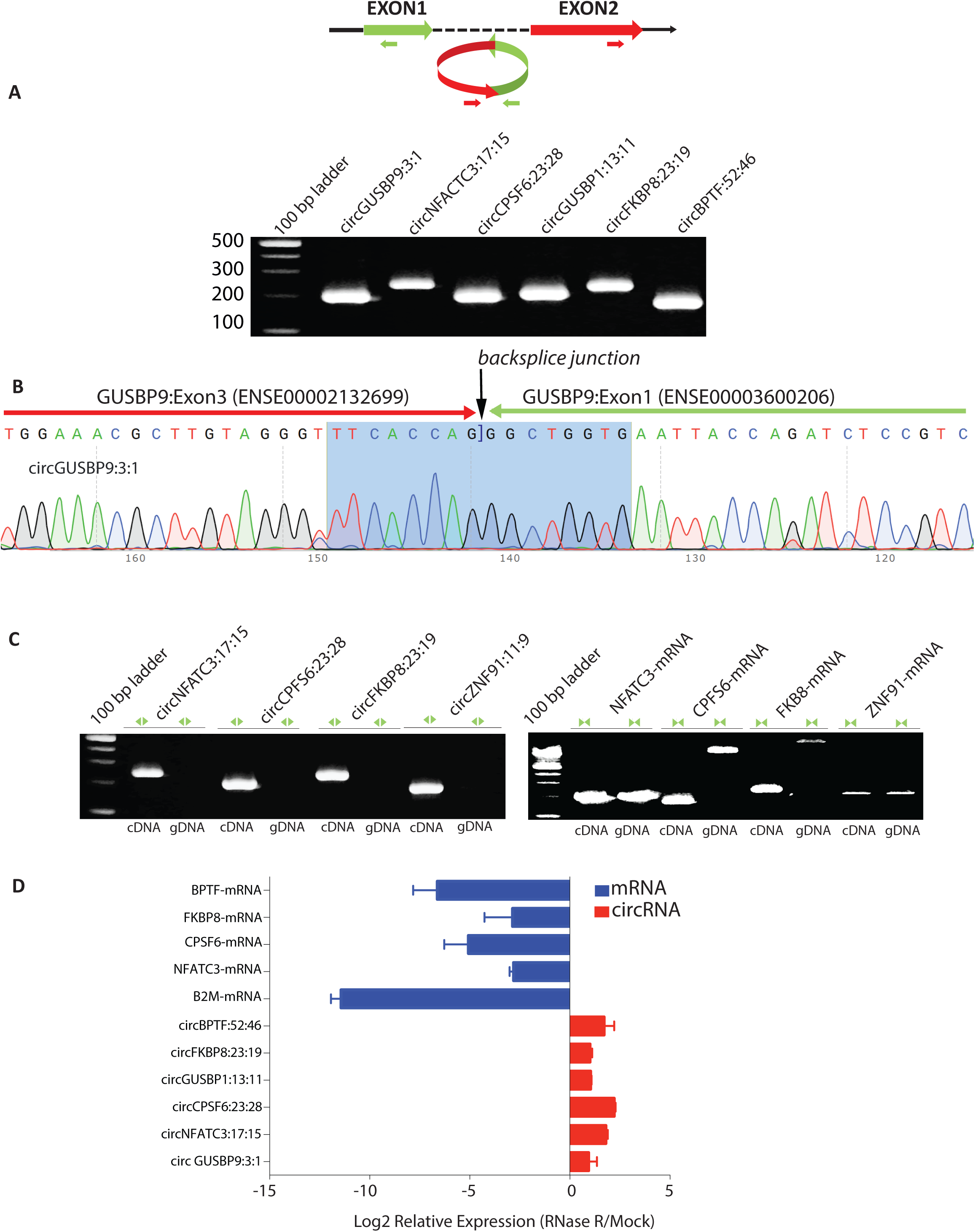
Validation of circRNA candidates through RT-PCR and Sanger sequencing. (A and Supplementary_Figure_1.pdf) Divergent primers with respect to genomic sequence were designed for 17 circRNA candidates. These become properly inward facing and identify the backsplice junction sequence. (B and Supplementary_Figure_1.pdf) Sanger sequencing confirms the backsplice junction sequence of the PCR products. Arrows indicate the presence of backsplice junction. (C) Divergent primers with respect to genomic sequence amplify circRNA backsplice junction sequence in cDNA but not in gDNA fraction. Convergent primers with respect to genomic sequence amplify mRNA in both cDNA and gDNA fractions. (D) Treatment with RNase R leads to enrichment of circRNAs and depletion of host gene and Beta-2 microglobulin (B2M) mRNAs compared to mock.

### Assessing circRNA associated genomic variants

Genotype calls corresponding to the 358 LCL samples were extracted from VCF files downloaded from Phase 3 release of the 1000 Genomes project [27]. Only common single nucleotide polymorphic (SNP) variants with a minimum allele frequency of >5% and passing Hardy-Weinberg Equilibrium filter were taken for association testing. The significance of correlations between genotypes and circRNA expression levels were determined by linear regression on quantile normalized circRNA expression values, after correction for known and inferred technical covariates (see methods), using Matrix eQTL [33]. The results of this additive linear genetic model association analysis are summarized in Figure 2. We tested ∼6.2 million SNPs and 95,675 circRNAs for cis QTL associations (within 1MB of gene boundaries) and plotted the genome-wide distribution of the test statistic against the expected null distribution to observe any inflation or residual deviations in the association statistic that may lead to excessive false positive rate. The observed distribution of p-values deviates from the expected null distribution in the extreme right tail only indicating no evidence for systematic spurious associations (Figure 2A). The nominal p-values obtained from Matrix-eQTL were adjusted for multiple testing using a linkage disequilibrium aware method [34]. We defined “ecircRNAs” as circRNAs with at least one SNP in cis significantly associated (adjusted p-value of < 0.05) with expression of that circRNA. A genome-wide p-value limit obtained as the median of the p-values for the best circQTL in each gene is used as a cutoff to identify all the circQTLs. This resulted in a total of 139,485 circQTLs for 2260 ecircRNAs in 1359 genes (Supplementary_File_S4.xlsx). The circRNA expression level of these genes does not seem to significantly influence the number of identified circQTLs (Supplementary_Figure_2.pdf). We also observed an enrichment of the circQTL variants at the proximity of the backsplice junction (Figure 2C). As mapping of reads to measure the circRNA expression could show allelic biases due to sequencing polymorphisms within the backsplice forming exons and hence could act as a cofounder for the association signal [35]. Thus, if one allelic read maps better to a location than compared to the other allelic read, a strong false positive association would be induced, generally closer to the measured event. We found only 721 out of 139485 circQTLs encompass the backsplicing exons, strongly indicating absence of any allele-specific read mapping biases in our results.

**Figure 2.**
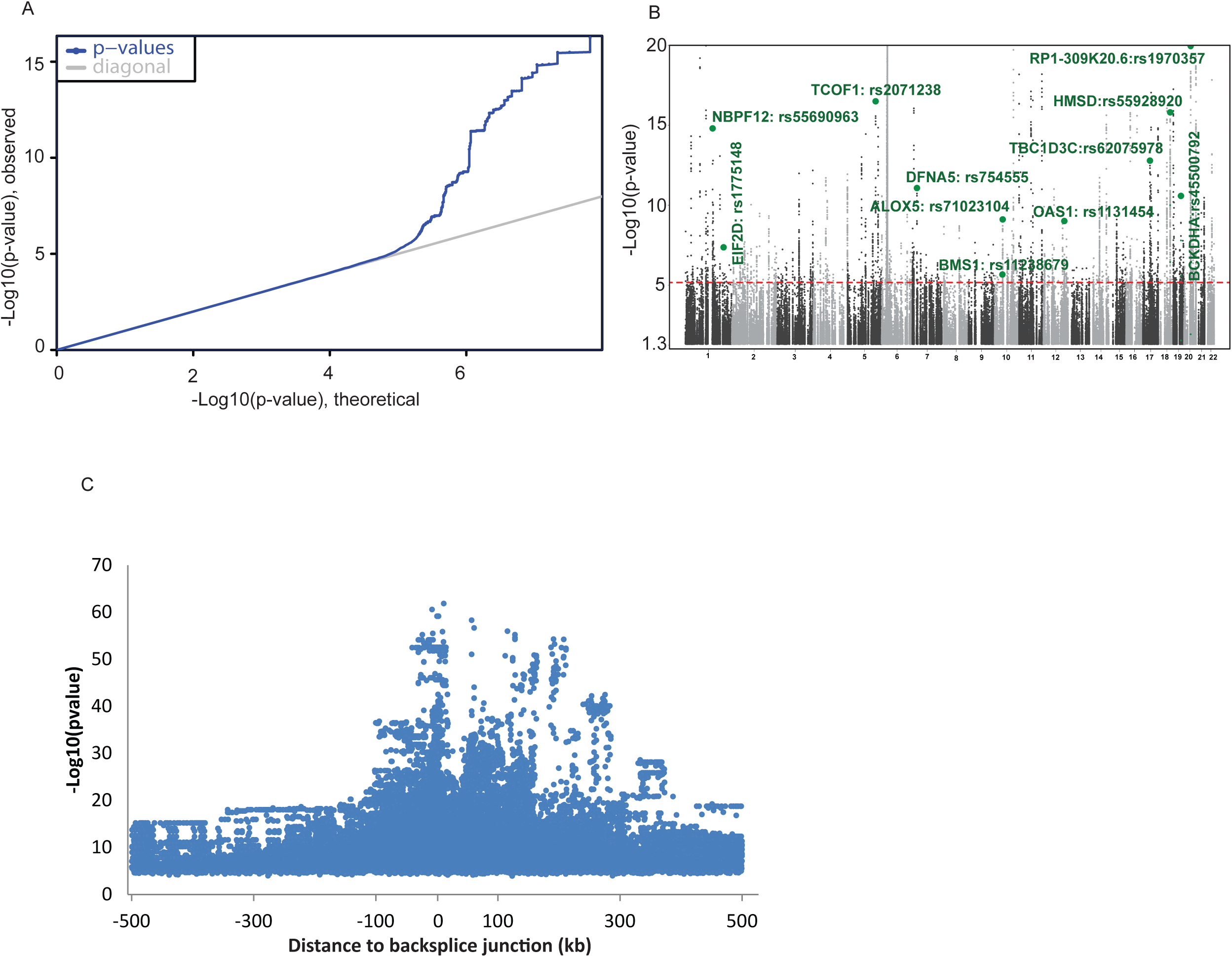
(A) Q-Q plot distribution of all recorded p-values for chromosome 22. (B) Manhattan plot of association statistics. The -log 10 (p-value) is plotted against the physical positions of each SNP on each chromosome and have been capped to P = 1E-20. The genome-wide significance threshold of 8.4E-06, obtained as the median of the unadjusted p-values for the best circQTL of each ecircRNA is shown as a horizontal red line. (c) The distribution of statistical significance of association for circQTL SNPs against proximity to circular RNA.

Many of the genes containing ecircRNAs, are associated with a disease phenotype (ALOX5, TCOF1, DFNA5, OAS1, EIF2D) or a physical trait like BMI and Height (BMS1, RP1-309K20.6; Figure 2B). For example, the gene Arachidonate 5-Lipoxygenase (ALOX5), has been associated with chronic bronchial asthma susceptibility and we found several circQTL SNPs in the promotor of ALOX5 gene that are associated with the expression of circular RNA processed from exons 1 and 2 of this gene (Figure 3).

**Figure 3:**
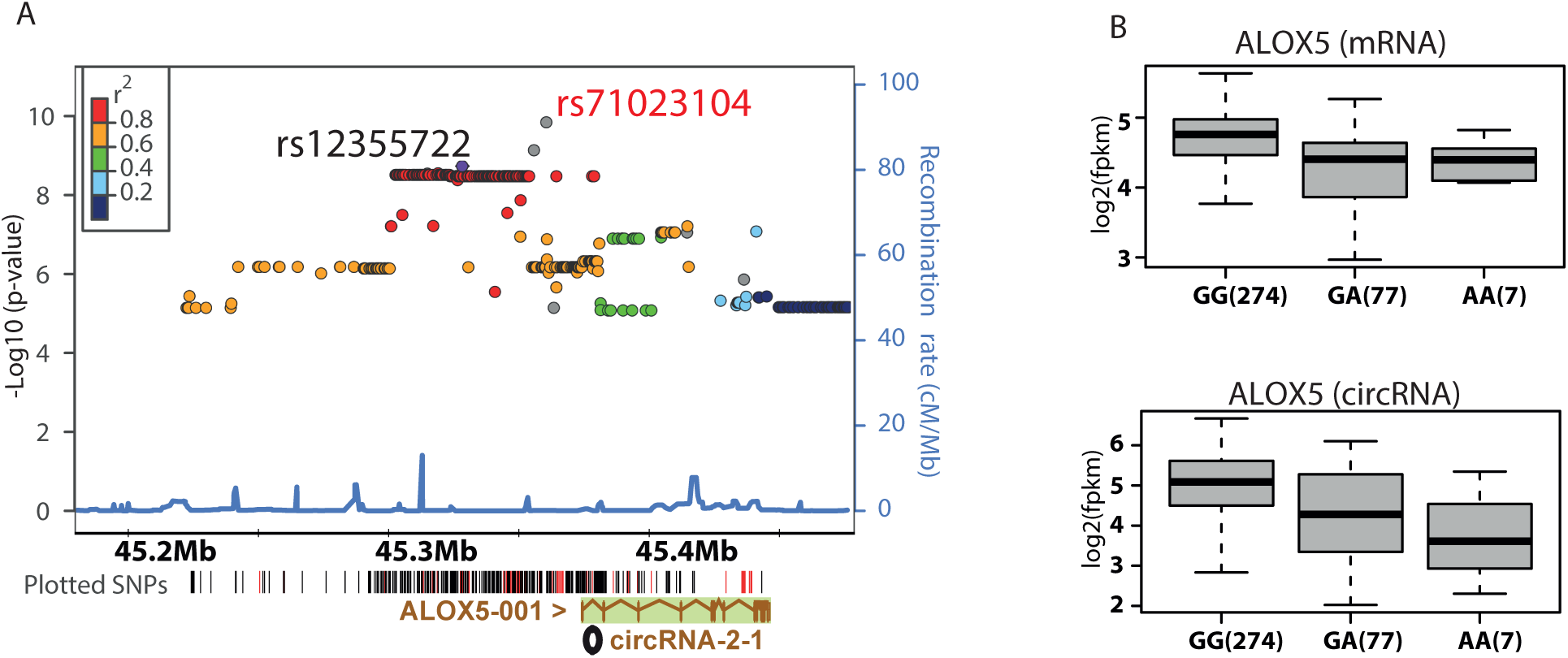
(A) Region association plot for ALOX5 circRNA. The plot is centered at the locus rs12355722 (purple diamond), which is associated with both circRNA and mRNA expression of ALOX5 gene. The circQTL only SNP (rs71023104) shows highest association with ALOX5 circRNA processed from its exons 1 and 2. In the plotted SNPs, red color indicates variants associated with only circRNA expression and black shows variants associated with both circRNA and mRNA expression. The values of r^2^ are based on the CEU HapMap 2 samples. The CEU HapMap 2 recombination rates are indicated in blue on the right y-axes as obtained using LocusZoom (http://csg.sph.umich.edu/locuszoom). (**B**) The circQTL and eQTL variant rs12355722 is associated with both circRNA and mRNA expression of the gene and shows a higher effect size beta (circRNA:mRNA −0.47:−0.35) for circRNA. Number of samples in each genotype group is shown in the brackets.

### circRNA QTLs exist independently of eQTLs

We compared our results for ecircRNAs with that of eQTL associations obtained by Lappalainen et al. in Geuvadis project [24]. Out of a total of 1359 genes containing ecircRNAs, 75% of these genes have no corresponding eQTLs in the Geuvadis data and ∼62% of the genes with only eQTLs have no evidence for circRNA expression. Only one quarter of the ecircRNA genes have one or more eQTLs associated with the linear expression of the gene (Figure 4A). Of these genes, that contain both circQTL and eQTL SNP markers, only ∼7% of the QTLs are shared having significant associations with both the circRNA and mRNA expression (Figure 4B). Thus, in these genes 74% of the circQTL markers and 90% of the eQTL markers are exclusively associated with expression of their respective isoform. In order to determine whether circQTLs are associated with any measurable effect on mRNA expression of the host gene, we compared the circRNA and mRNA expression changes between homozygous (REF) and heterozygous genotypes of circQTL markers. Figure 4C shows circRNA and mRNA expression changes between homozygous and heterozygous genotypes for the best circQTL SNP of 1026 ecircRNA genes that have no corresponding eQTL. Though circRNA expression shows a clear trend for both positive and negative beta values (effect size), there is no such trend in the mRNA expression. Similarly, in Figure 4D, there is no significant difference in mRNA expression between homozygous (REF) and heterozygous genotypes for the 16205 circRNA exclusively associated SNPs in genes that contain both circQTL and eQTL SNPs. Thus circQTL markers do not seem to have any significantly measurable effect on the mRNA expression of their host genes, ruling out any artificial increase in spurious associations for ecircRNAs due to systematic or analytical biases from mRNA. These results also imply that QTL SNP markers exclusively associate with either circRNA or mRNA expression. Thus, a large fraction of ecircRNA genes exist without eQTLs, indicating the lack of total biological dependence of circRNA expression on eQTLs.

**Figure 4.**
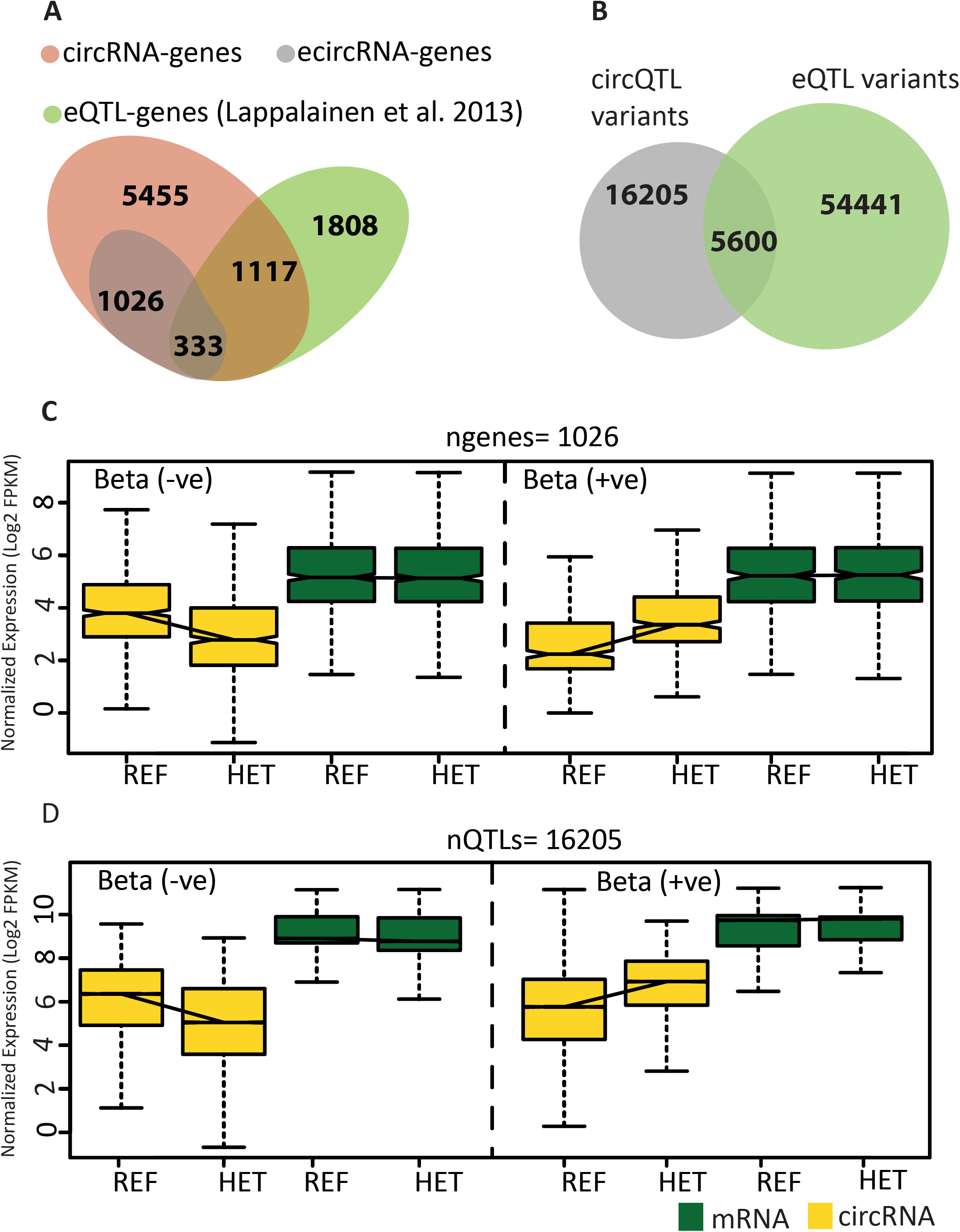
(A) Venn diagram showing overlap of ecircRNA genes with eQTL genes from Lappalainen et al. (**B**) Venn diagram showing distribution of genomic variants associated with circRNA and mRNA expression in 333 genes that contain both circQTL and eQTL SNPs. Only ∼7% (5600) QTLs are shared and show association with both circRNA and mRNA expression in these genes. (**C**) Boxplots representing distribution of average expression from all individuals with a given genotype. The trends in circRNA and mRNA expression between reference homozygous and heterozygous genotypes for best circQTL SNP, in 1026 ecircRNA genes that do not contain any eQTL associations. (**D**) circRNA and mRNA expression trends for 16205 circQTL SNPs from genes having both circQTL and eQTL markers.

### Shared QTLs have similar effects on circRNA and mRNA expression

To understand the effect of shared QTLs, we plotted circRNA and mRNA expression profiles for 5600 QTL markers that were significantly cis-associated with both circRNA and mRNA expression (Figure 5A). The trends in expression from homozygous to heterozygous genotypes are analogous for circRNA and mRNA, as would be expected based on co-transcription. Figure 5B, further evaluates the effect of the shared QTLs, by charting beta values of circQTL and eQTL associations in a scatterplot. The effect size (beta) defined as the slope of the linear regression indicates direction of change in expression from reference to alternate allele with negative values implying higher expression for the reference allele.

**Figure 5.**
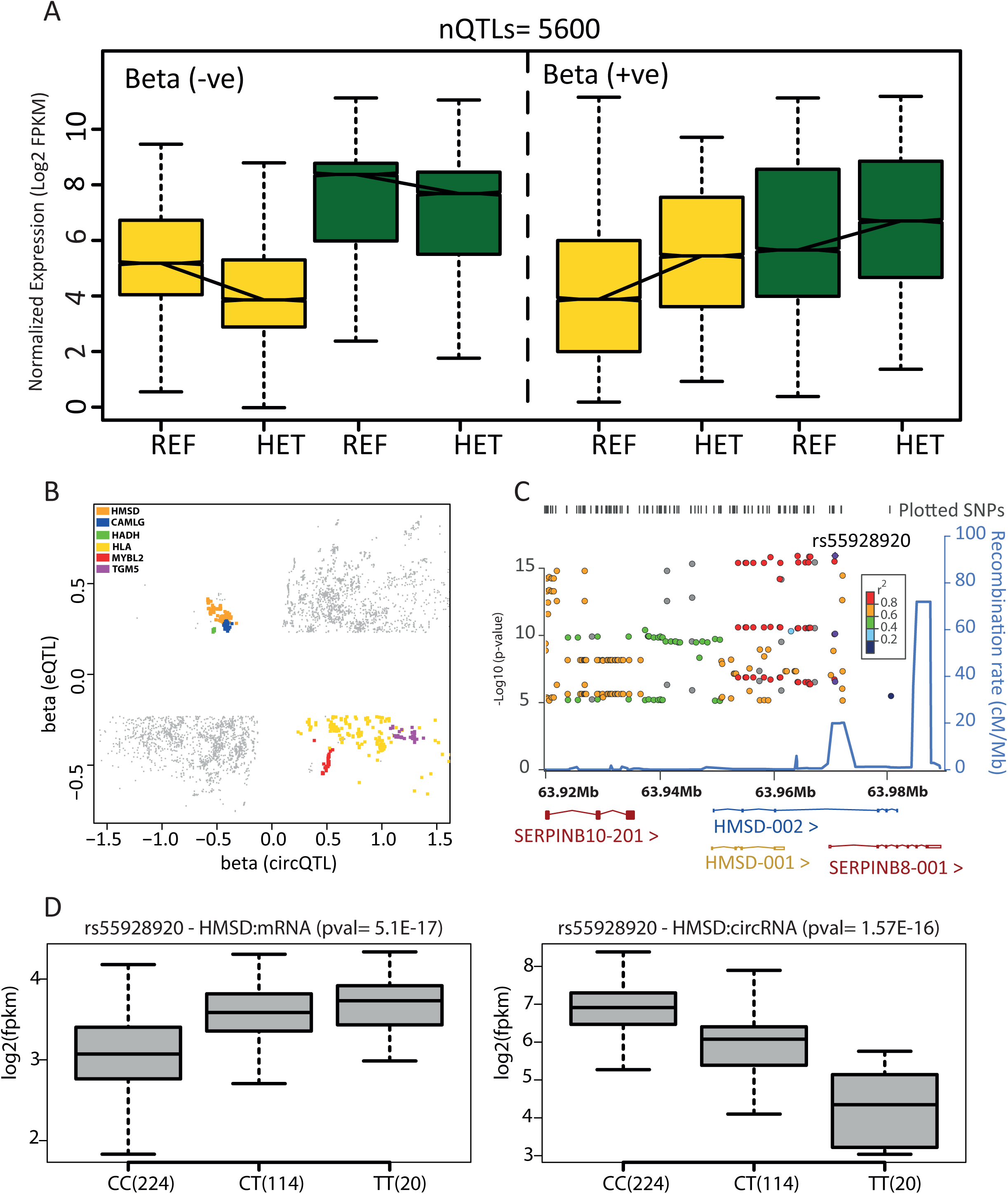
(A) Boxplots representing distribution of average circRNA and mRNA expression from all individuals with a given genotype for shared QTLs. (B) Spearman rank correlation defines the effect size (beta) and indicates direction of change from reference to alternate allele. Each point in the plot is a shared QTL marker with x-axis representing beta for circRNA association and y-axis representing beta for mRNA association. The high density of points in the upper right and bottom left quadrants show that most shared QTLs have similar effects from their reference and alternate alleles on circular and linear expression. Colored points represent QTLs that have opposite effects on circular and linear expression of their host genes. (C) Region association plot for HMSD circRNA. The shared intronic QTL (rs55928920) is associated with HMSD circRNA (pval= 1.57E-16) as well as mRNA expression (pval=5.1E-17). Values of r2 are based on the CEU HapMap 2 samples. CEU HapMap 2 recombination rates are indicated in blue on the right y-axes as obtained using LocusZoom (http://csg.sph.umich.edu/locuszoom). (D) Reference allele of rs55928920 is associated with higher expression of circRNA from 2^nd^ and 4^th^ exons of the HMSD transcript (HMSD-002) and lower expression of the mRNA transcripts from HMSD and SERPINB8 genes.

Though most shared QTLs have similar effects, a small subset has opposite effects on the circular and linear expression of their host gene. For example, the Histocompatibility Minor Serpin Domain Containing (HMSD) gene contains a cluster of 306 QTL markers that are associated with the expression of its linear transcript as well as a circRNA formed by the 2nd and 4th exons of the HMSD-002 transcript (Figure 5C). These shared QTL markers seem to drive the circular and linear expression of the HMSD gene in opposite directions, suggesting potential modulation of the circRNA levels at the expense of mRNA. The reference allele of the most highly associated circQTL marker (rs55928920) alleviates the expression of the HMSD circRNA while the alternate allele favors mRNA expression of HMSD and SERPINB8 genes (Figure 5D). We tested whether genes with shared QTLs having opposite effects on circRNA and mRNA abundance levels are expressed at similar scales in comparison to other ecircRNA genes (Supplementary_Figure_3.pdf) – the results show no difference in mRNA expression but a statistically significant higher circRNA expression of these genes compared to other ecircRNA genes.

### Sharp LD breakdown between circQTL and eQTL SNPs

Linkage disequilibrium (LD) is the non-random association between alleles at different loci, and can be estimated using the correlation between SNP markers [36]. To assess if circQTL and eQTL SNPs are largely independent, we asked how often a top eQTL SNP is also a circQTL SNP and vice-versa. We found only 43 instances where a top eQTL SNP was also a circQTL SNP for the gene and 72 instances where a top circQTL SNP was also an eQTL SNP for the gene. We further explored the extent of LD structure between circQTL and eQTL markers by computing the average pairwise correlation for shared QTL, circQTL and eQTL SNPs and compared these distributions with a cross-pair between circQTL and eQTL markers. Figure 6A indicates that eQTL SNPs tend to encompass weaker LD regions compared to circQTL and shared QTL markers. There is a significant decrease in the average pairwise correlation between a circQTL and eQTL cross-pair, when compared to intra-pairwise correlation distributions of shared QTL, circQTL and eQTL markers (median pairwise correlation drops from 0.8 to 0.3). This strongly suggests a sharp breakdown in the LD structure between circQTL and eQTL SNP containing regions. Sharp LD breakdown between circQTL and eQTL variants could indicate differential selection pressures shaping their molecular evolution.

**Figure 6.**
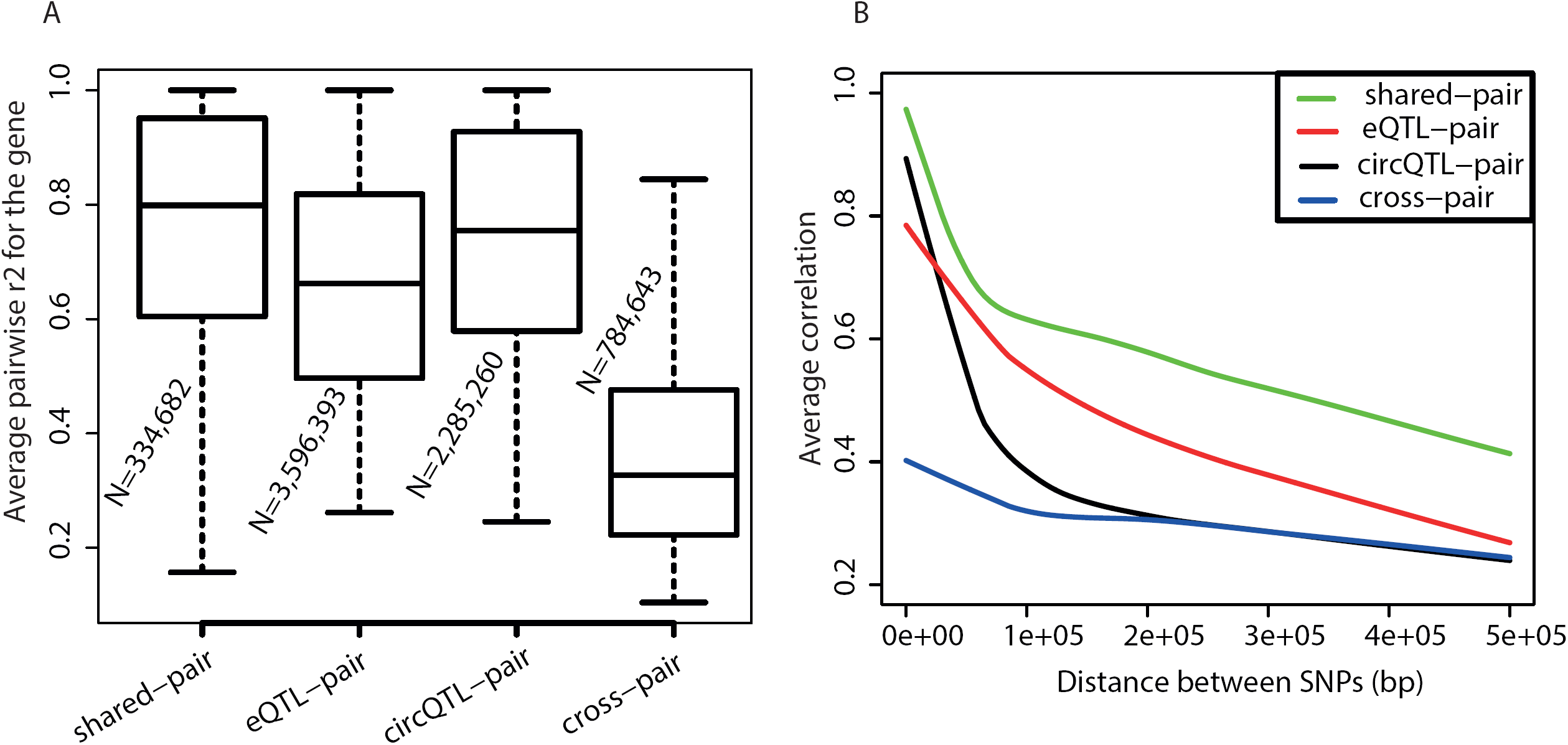
(A) Boxplot distributions of the average pairwise correlation (r^2^) between shared QTL, eQTL, circQTL and cross-QTL SNPs for each gene. Average pairwise correlations were computed for each category of QTL markers for genes containing both circQTL and eQTL SNPs. Cross-pair is the pairwise correlation between a circQTL and an eQTL marker. For each boxplot distribution, N represents total number of pairwise combinations from all genes; only combinations with r2 above 0.1 were used for the analysis. (B) Loess fit curves for average correlation (r2) vs distance between the QTL markers.

To estimate the rate of LD decay for circQTL and eQTL markers and compare their LD patterns, we plotted the loess curves of average pairwise correlations against the physical distance between the SNP pair (Figure 6B). LD for circQTL SNPs decayed faster than eQTL and shared QTL markers. Average pairwise correlation for circQTL variants drops sharply within a span of 50kb, indicating that circQTLs are held in tightly linked and comparatively shorter LD blocks. For the cross-pair between circQTL and eQTL SNPs, LD shows very minimal change with distance, indicating the lack of genetic linkage between the two sets of these markers.

### Functional characterization of circQTL variants

In order to assess the functional elements of the genome enriched for circQTL variants, we performed Monte Carlo simulations using 10,000 random sets of background eQTL SNPs with minor allele frequencies (MAF) matched to the set of 23769 circQTL variants (see Methods). By functionally classifying the circQTL and background eQTL SNPs as per the definitions in SnpEff [37], we found that circQTLs are significantly enriched among variants in 5’-untranslated (Monte Carlo *P-value 2.3E-02*), exonic (Monte Carlo *P-value* 8.61E-03), intergenic (Monte Carlo *P-value < 1E-04*) and regulatory (Monte Carlo *P-value 9.94E-04*) regions of the genome. In contrast, surprisingly, there is a significant underrepresentation of circQTL SNPs among intronic region variants (Monte Carlo *P-value < 1E-04*) (Figure 7 A,B). Though circQTL variants were also found to be enriched for splice site regions (odds ratio (OR)=1.55), the *P-value* for this comparison was insignificant (*P = 1.99E-01*), possibly due to a low fraction of variants annotated as “splicing”.

**Figure 7.**
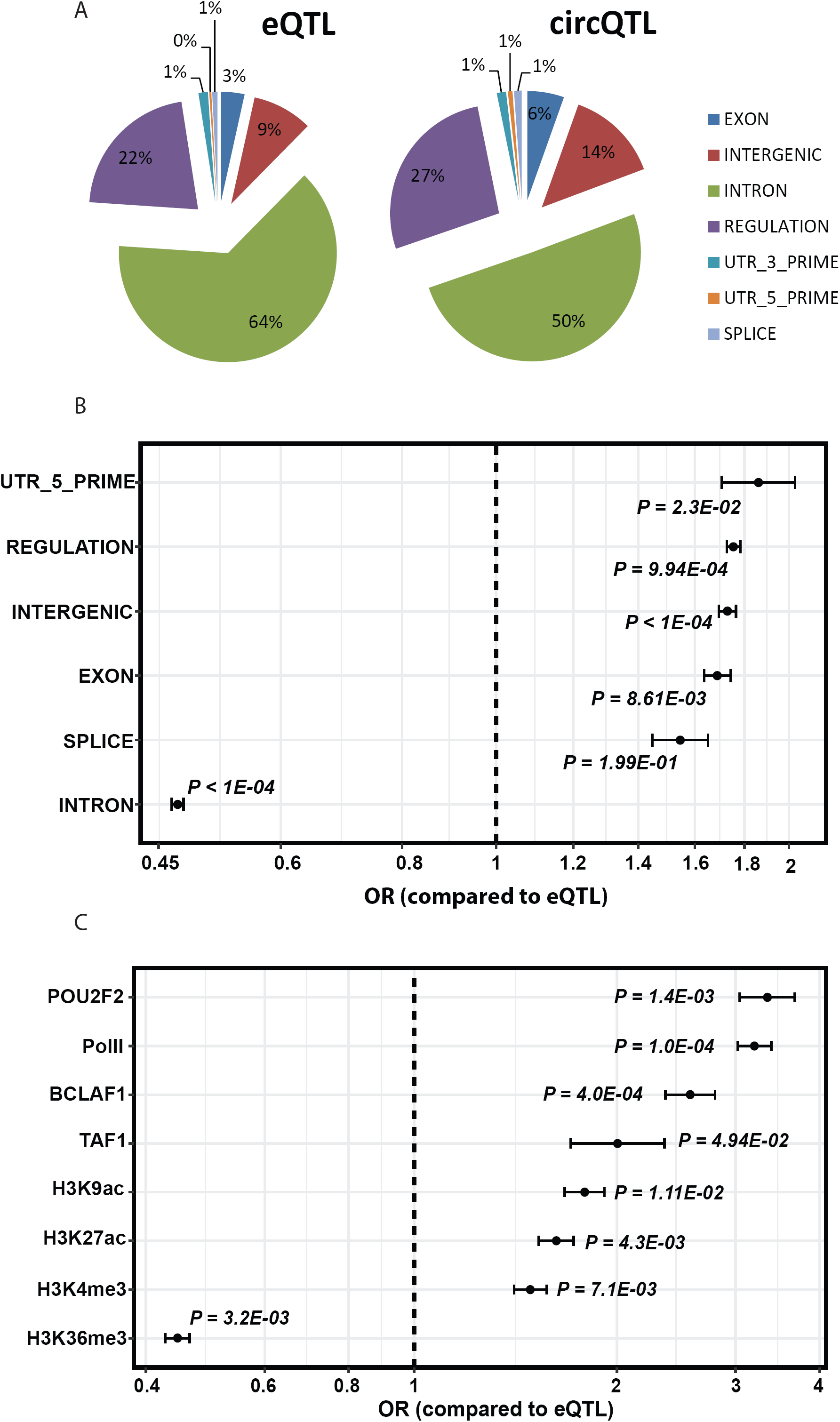
(A) Proportion of SNPs annotated for each functional category using SnpEff [37]. (B) Odds ratio (OR) for each functional category compared to the eQTL background. (C) Enrichment of circQTL variants within genetic regulatory elements. OR greater than 1 indicates enrichment of circQTL variants. The significance of the OR was tested using 10,000 Monte Carlo permutations of the eQTL SNPs, and the significance level is reported as a Monte Carlo P-value. Bars indicate 95% confidence intervals.

To further explore if chromatin structure has a perceivable influence on circRNA expression, we analyzed the enrichment of circQTL variants among various genetic regulatory element sequences. These regulatory sequences correspond to cell-type specific tracks of chromatin modifications, RNA polymerase (Pol II, Pol III) bound regions and transcription factor binding sites for GM12878 (lymphoblastoid) cell line obtained from the ENCODE project [38], and used to annotate circQTL and background eQTL variants. Using Monte carlo simulations, 10^4^ iterations were performed to annotate a randomly generated set of background eQTL SNPs with MAF matched to circQTL SNPs and the odds ratio and significance of the enrichment computed for each chromatin mark. A total of 42 regulatory marks were analyzed and significant enrichment of circQTL SNPs was found among variants contained in transcriptionally active chromatin regions. Histone modifications associated with active transcription that showed most significant enrichments were acetylated histone H3 lysine 9 (H3K9ac; OR = 1.79; P = 1.1E-02), acetylated histone H3 lysine 27 (H3K27ac; OR = 1.63; P = 4.3E-03) and trimethylated histone H3 lysine 4 (H3K4me3; OR = 1.49; P = 7.1E-03). On the contrary, a significant depletion of circQTL SNPs was observed among trimethylated histone H3 lysine 36 (H3K36me3; OR = 0.44; P = 3.2E-03) marked genomic regions. We also found significant enrichment of circQTL variants for Pol II RNA polymerase bound regions and POU2F2, BCLAF1 and TAF1 transcription factor binding sites (Figure 7C).

The potential of circQTL SNPs in context of their contribution to disease risk and phenotype association was analyzed using the NHGRI-EBI GWAS Catalog [39], which is a curated collection of all the published genome-wide association studies for various human diseases and traits. We tested the enrichment of circQTL SNPs for various Experimental Factor Ontology [40] (EFO) terms defined in the catalog and representing a disease or a trait. The circQTL and a random set of background eQTL SNPs of matched MAF were separately grouped into genomic loci based on their LD patterns, and defined as the genomic region containing SNPs in LD with the index SNP at r^2^ > 0.8. An associated genomic locus for a trait is identified if a GWAS risk SNP falls within the locus. Finally, the enrichment of circQTL SNPs among associated loci is evaluated by a one-tailed Fisher’s exact test followed by Bonferroni correction for the total number of traits assessed. The top diseases associated with the genomic loci containing a significant enrichment for circQTL SNPs are schizophrenia (OR = 4.06, P = 2.02E-23, one-tailed Fisher’s exact test with Bonferroni correction), nephropathy (OR = 63.6, P= 7.08E-08), plantar warts (OR = 50.7, P= 8.27E-06), disc degeneration (OR = 14, P = 3.4E-03), lung cancer (OR = 4.24, P = 3.49E-03) and Asthma (OR = 8.45, P = 4.49E-03) (Figure 8). It has been proposed that QTLs associated with the alternative splicing events could causally contribute to the risk of schizophrenia[25], and hence circQTL SNPs having the most significant enrichment for schizophrenia could provide a vital resource for assessment of the disease risk or uncovering of the disease causality from cis-acting genetic variants.

**Figure 8.**
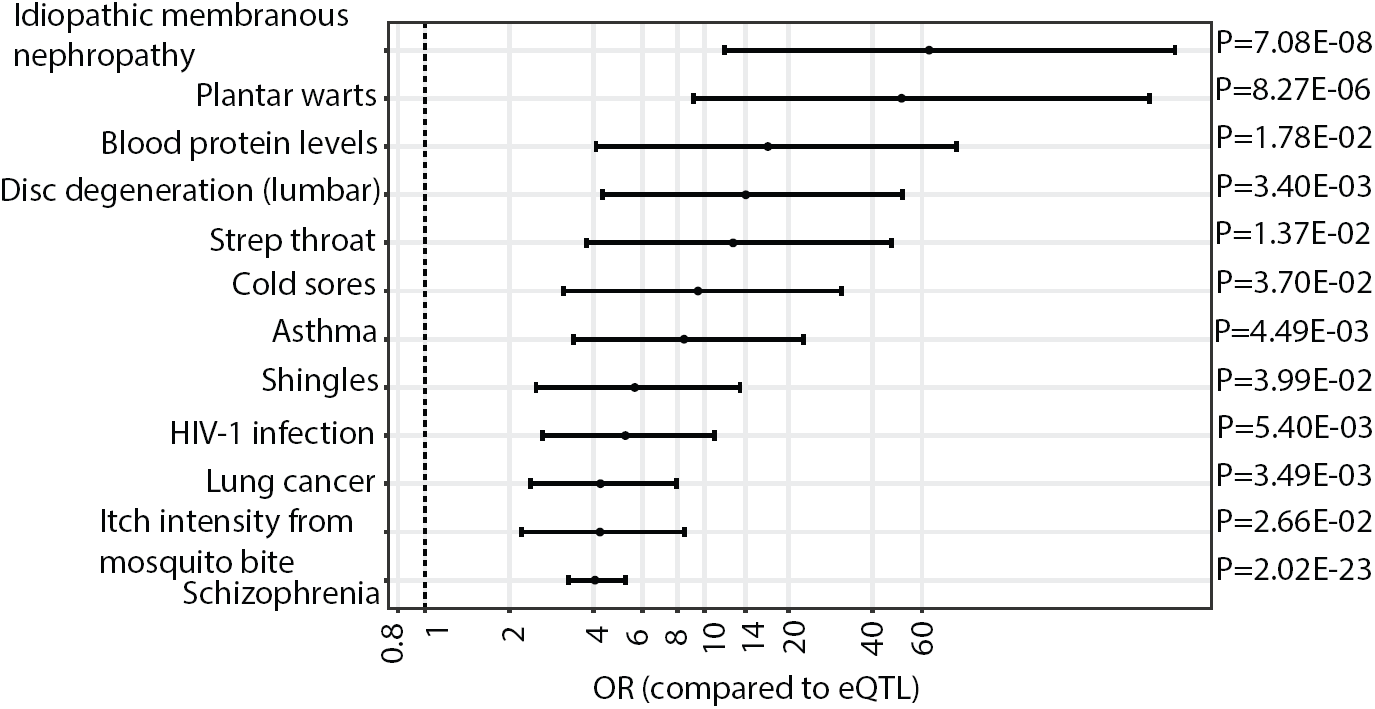
Enrichment of circQTL SNPs among disease associated loci. Results of enrichment analysis of circQTL SNPs associated with various traits in GWAS catalog [39] and arranged by odds ratio. A total of 306 traits were tested and only 12 were found to show statistically significant enrichments. The nominal P-values obtained from one-tailed Fisher’s exact test were Bonferroni corrected for the total number of 306 traits tested and a p-value cut-off of 1.6E-04 (0.05/306) was used as a significance threshold. Bars indicate 95% confidence intervals.

## Discussion

In this study we quantified the contribution of cis-acting genetic variants on circular RNA expression from RNA-Seq data of the 1000 Genomes project lymphoblastoid cell lines [24]. To our knowledge, this is the first study to comprehensively identify the set of regulatory variants influencing the circRNA expression in a single cell type.

Using our circRNA detection pipeline [28], we identified 95,675 circular RNAs which are likely to be adequately expressed in human peripheral blood transcriptomes and thus constitute an important resource. Our genome-wide analysis for association of circRNA expression with genotype identified thousands of previously uncharacterized cis-acting genetic variants influencing circRNA transcript abundance. We identified a total of 1359 genes with circQTL associations, referred here as ecircRNA genes. These ecircRNA genes are enriched in canonical pathways controlling important cellular processes like cell cycle regulation and spliceosome formation. Many of the ecircRNA genes are linked to a disease phenotype (ALOX5, TCOF1, DFNA5, OAS1, EIF2D) or a physical trait like BMI and Height (BMS1, RP1-309K20.6). The gene Arachidonate 5-Lipoxygenase (ALOX5), which contains several circQTL SNPs in its promotor region has been linked to chronic bronchial asthma susceptibility and encodes a member of the lipoxygenase gene family that catalyzes the conversion of arachidonic acid to leukotriene A4.

Leukotrienes are important mediators of a number of inflammatory and allergic conditions and mutations in the promoter region of this gene have been shown to lead to a diminished response to anti-leukotriene drugs used in the treatment of asthma [41]. The presence of several upstream circQTL SNPs exclusively associating with the circRNA transcript processed from exons 1 and 2 of this gene could indicate causal connection of the circQTLs with drug response, acting thereby through modulation of circRNA levels.

By evaluating genes for the presence of circQTLs or eQTLs and the influence of these SNPs on host gene expression, we were able to show that most QTL SNP markers exclusively associate with either circRNA or mRNA expression. A large fraction of ecircRNA genes do not have eQTLs, indicating an arrangement at the level of DNA sequence for circRNA regulation that does not seem to influence the basal gene expression. Likewise, circQTL markers do not have any significantly measurable effect on the mRNA expression of their host genes reinforcing the idea that circRNA and mRNA transcription could involve distinctive regulatory mechanisms.

In genes that contain both circQTLs and eQTLs, only a small fraction of the markers are shared between the two associations. These shared QTL markers frequently, but not always, have similar effects on the linear and circular RNA expressions of the gene. Some shared QTL markers, however, have opposite effects on gene expression potentially driving the circRNA levels at the expense of mRNA.

We compared the LD patterns for circQTL and eQTL markers and found that eQTL SNPs tend to encompass genomic regions of weaker LD structure compared to circQTL and shared QTL SNPs. The circQTL SNPs are held in tightly linked LD blocks with their LD structure decaying faster than eQTL and shared QTL markers. A sharp breakdown in the LD structure exists between circQTL and eQTL SNP containing genomic regions. Thus differential selection pressures could be working on circQTL and eQTL SNPs to influence their distinct regulatory mechanisms on gene expression.

We further characterized the circQTL SNPs for relationship to various annotated genomic features. We found an enrichment of circQTL SNPs among transcribed sequence variants including 5’-untranslated, exonic, intergenic and regulatory regions of the genome. The enrichment of circQTL SNPs in coding sequences is in accordance with the notion that coding sequences in complex genomes can simultaneously encode amino acid and regulatory information including the potential to act as transcriptional enhancers or splicing signals [42]. Coding exons have also been shown to function as tissue-specific enhancers of nearby genes operating as coding exons in one tissue and enhancers of nearby gene(s) in another tissue [43]. In this context, circQTL SNPs in coding exons could have dual functional roles with ability to either encode a transcript variant of its host gene or adjust a transcriptional regulatory signal for a nearby gene to alter its circRNA levels.

We also found that circQTL SNPs are enriched among genomic regions marked by chromatin modifications associated with active transcription. The enrichment of circQTL SNPs among histone modifications and transcription factors associated with an enhanced transcriptional activity could indicate a role for chromatin structure in regulating circRNA expression - possibly through aiding splicing machinery in coordinating splicing reactions. Several studies have suggested a cross talk between transcription and pre-mRNA splicing emphasizing a functional connection between RNA Polymerase II (RNAP II) elongation and the splicing machinery [44–47]. There is accumulating evidence that both RNAP II and the landscape of the chromatin marks contribute to regulate the splicing patterns through complex regulatory network that contains both feedforward and feedback loops. The kinetic model of transcription-coupled splicing regulation proposed by Kornblihtt et al. states that chromatin structure influences the rate of RNAP II transcription elongation, and the rate of transcription elongation in turn influences alternative splicing decisions [48,49]. Weak exons lead to RNAP II pausing and increased inclusion of the exon in spliced mRNA whereas increased elongation rate decreases exon inclusion in mRNA [46]. The chromatin modifications H3K4me3, H3K9ac and H3K27ac which show an enrichment for circQTL SNPs and mark active gene promotors and enhancers are known to play a role in alternative splicing through modulating the rate of RNAP II elongation [46,47,50,51]. On the other hand circQTL SNPs are underrepresented for H3K36me3, another histone mark characteristic of active transcription particularly from lowly expressed genes, but known to cause hypoacetylation and exon inclusions in the mRNA, possibly through inducing RNAP II pausing [47,52]. Therefore chromatin modifications, probably through a combinatorial pattern could provide RNAP II with signals that locally regulate elongation rate and alter the splicing decisions to favor either mRNA or circRNA expression. In this scenario certain combinatorial patterns of active chromatin modifications could induce a high RNAP II elongation rate that would translate into exon exclusions from mRNA and possibly their retention in the circRNA form.

As genetic variants can influence patterns of chromatin marks [53], some of the circQTL SNPs could function by altering the epigenetic status of various histone modifications thereby enhancing the RNAP II elongation rate which will promote a higher circRNA production. Alternatively, circQTL SNPs can have a more direct role through recruitment of transcription or splicing factors to influence the transcription rate which in turn could alter the epigenetic state of the DNA template.

We observed significant enrichment of circQTL SNPs with strong to moderate associations with various human diseases indicating role for circQTL variants in genetic disease etiologies. Thus circQTL SNPs with mostly exclusive associations to circRNA expression could potentially act as causal variants or explain an additional part of the genetic architecture of the disease, linking GWAS phenotypes and diseases to putative target genes and regulatory elements. With the findings presented here, we provide a set of SNPs for future investigations of the role of circular RNAs in the genetic and molecular mechanisms underlying diseases and traits.

## Methods

### Cell Culture

Human T lymphocyte Jurkat cells were cultured in RPMI (Sigma, St. Louis, MO, USA) supplemented with 10% fetal bovine serum (Life technologies, USA).

### RNA isolation, RNase R treatment and DNA isolation

Total RNA from Jurkat cells was isolated by using RNAesay mini kit (Qiagen Valencia, CA USA). For RNase R treatment, 20 units of RNase R (Epicentre), 1 microliter of RNase R buffer and 1 unit/microliter murine Ribonuclease Inhibitor (New England Biolabs) were added to the 2 microgram of RNA samples and incubated at 37°C for 30 minutes. Total DNA was isolated from Jurkat cells using a DNeasy kit (Qiagen).

### Sanger sequencing and Realtime PCR

1 μg of total RNA was used for cDNA synthesis using the iScript select cDNA Synthesis Kit (Bio-Rad). Outward facing primers with respect to genomic sequence were designed to specifically amplify the backsplice junctions of representative circRNAs in a PCR assay. PCR reactions were performed for cDNA samples and genomic DNA using 28 cycles. PCR products were analyzed by 2.2% agarose gel electrophoresis and gel purified PCR products were directly sequenced to identify the gel product.

For Realtime PCR Fast Start Universal SYBR Green Master mix (Roche) was used to amplify the backsplice junctions using divergent primers, primers for mRNA were obtained from Primer3 software. Each Real-Time assay was done in triplicate on Step One Plus Real time PCR machine (Life Technologies, CA, USA).

### Processing of genotype data

The 1000 Genomes Project phase3 release of variant calls was downloaded from ftp://ftp.1000genomes.ebi.ac.uk/vol1/ftp/release/20130502/. The variant set contained genotype data for 2504 individuals from 26 populations. The vcf-subset function from vcftools [54] was used to extract the genotype data for 358 european samples from the downloaded VCF files. These corresponded to 89 CEU, 92 FIN, 86 GBR and 91 TSI samples. We recomputed the allele number and allele counts with -- recode --recode-INFO-all parameters for the defined number of samples and the variants with a resulting minimum allele frequency of >5% were taken for further analysis. Further variants that didn’t pass HWE test were also filtered out. The original VCF files were GRCh37 reference assembly mapped. We used the All.vcf.gz file downloaded from ftp://ftp.ncbi.nlm.nih.gov/snp/organisms/human_9606_b142_GRCh38/VCF/ and containing GRCh38 mapped coordinates for 1000 genome variants to remap the GRCh37 variant coordinates from original VCF files to GRCh38 cords using custom Perl scripts.

### Processing of RNA-Seq data

RNA-Seq data for 358 european individuals corresponding to our genotype samples was download from http://www.ebi.ac.uk/arrayexpress/experiments/E-GEUV-1/samples and its mapping with the genotype information was obtained from: http://www.ebi.ac.uk/arrayexpress/files/E-GEUV-1/E-GEUV-1.sdrf.txt. CircRNA pipeline [28] was run on all 358 RNA-Seq data sets on a Linux cluster managing batches of jobs as per the load on the cluster. Circular RNA structures are identified by using Bowtie 2 [29] to align paired-end RNA-seq data to a custom reference exome containing all possible pairs of intragenic non-linear combinations of exons, as well as single exon “backsplices”. A scrambled junction is inferred as a circRNA candidate when one mate of a paired-end read aligns at the backsplice junction with a minimum of 10bp overlap on either side of the junction and the other mate aligns in the body of the backsplice forming exon/s. Supportive evidence is also considered when mates of a paired-end read align to the exons in divergent orientation with respect to the genomic sequence, suggesting a scrambled junction instead of a linear junction. These alignments are further filtered to remove any potential PCR duplicates and only primary alignments are kept for final assessment of circRNA expression [28]. The expression of the circRNA in each sample is then defined as the sum of junctional read-pairs and supportive read-pairs. With all the 358 samples, aligned and circRNAs called, results were merged into one data table with rows as circRNA name and columns as counts from each sample. The pipeline initially reported 3,332,457 backsplice junctions with at least 1 read pair in 358 samples. We applied first filtering cutoff for a minimum of 50 junctional reads across the samples and this resulted in 323900 circRNA candidates. As we also consider supportive evidence, the minimum of 50 junctional read-pairs could come from either one sample or multiple samples. Nonetheless, the requirement for a high number of minimally seen junctional reads mostly translates to have the junctional evidence coming from multiple samples rather than a single sample. A further filtering for expression in at least 60% of the samples and outliers removed (expression < 500 or greater than 50000 counts) resulted in 95675 circRNAs that were taken for the final association analysis.

### circQTL calling

We used Matrix eQTL [33] to obtain the significance of the correlation between genotypes and circRNA expression after correcting for known and inferred technical covariates. The genotype VCF files were converted into Matrix_eQTL input genotype and snploc files using custom Perl scripts. The raw quantifications for circRNA expression were library size and length corrected (converted to fpkm), quantile normalized and log2 transformed and association with genotype was determined by linear regression on normalized circRNA expression values. For controlling potential confounding factors, we included first three principal components from the genotype data, RNA library sizes, population ID and RNA sequencing lab ID as covariates in the regression model. All SNPs that lie within 1Mb region of the gene boundaries were tested for cisQTL associations with the circRNA expression of the gene. The p-values obtained from Matrix-eQTL were adjusted for multiple testing using eigenMT package, which uses a linkage disequilibrium aware method for multiple testing correction in eQTL studies [34].

### Matched control sets

The circQTL SNPs that are exclusive associated with the circRNA expression were used for all enrichment analysis (functional/regulatory/disease). The total number of 139485 circQTLs identified in this study, correspond to 23769 unique SNPs exclusively associated with circRNA expression. We used this list to generate BED files of random background sets of eQTL SNPs for comparing the enrichments. First the set of eQTL SNPs from [24] was filtered using a custom script to only include SNPs which are not in LD with the circQTL SNPs (r2 < 0.2). Finally, using circQTL and non-LD eQTL lists of SNPs as input in UES [39], we generated random background sets of eQTL SNPs for use in enrichment analysis.

### Functional annotation enrichment analysis of circQTL SNPs

To conduct functional enrichment annotation analysis for circQTL SNPs, we generated 10,000 random background datasets of matching eQTL SNPs using the process described above. The random sets generated by UES [39] are in BED file format. We extracted the SNP IDs from BED files and used them in vcftools [54] to generate VCF files for the background data. Similarly, using vcftools, a VCF file was generated for the circQTL SNPs. We used SnpEff [37], with the options (-csvStats -spliceSiteSize 20 -ud 0 -no-intergenic -no-utr -v –reg) to annotate both circQTL and the eQTL background sets of VCF files for the following functional annotation terms defined for GRCh38.86: EXON, INTERGENIC, INTRON, REGULATION, SPLICE_SITE_ACCEPTOR, SPLICE_SITE_DONOR, SPLICE_SITE_REGION, UTR_3_PRIME, UTR_5_PRIME. The output csv files generated by SnpEff are processed using custom scripts to compute odds ratio (OR), 95% CI and p-value of the enrichment. The results of the SPLICE_SITE_ACCEPTOR, SPLICE_SITE_DONOR, and SPLICE_SITE_REGION terms were combined and reported as “SPLICE”. The Monte Carlo P-value for the enrichment or underrepresentation of each functional term was calculated as follows:

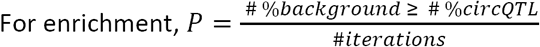

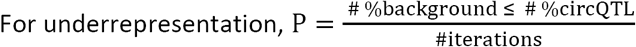

Where,

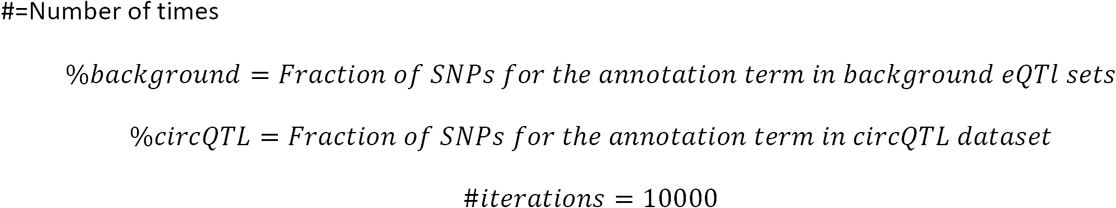

### Regulatory elements enrichment analysis of circQTL SNPs

Encode annotations of the regulatory tracks corresponding to GRCh38.86 was downloaded through SnpEff using its download option. We generated 10,000 random background datasets of matching eQTL SNPs using the process described above and converted them into VCF files as described in the functional annotation enrichment analysis. We used SnpEff [37], to annotate both circQTL and the eQTL background sets of VCF files for these regulatory tracks of lymphoblastoid (GM12878) cell line: ATF3-GM12878_enriched_sites BATF-GM12878_enriched_sites BCL11A-GM12878_enriched_sites BCL3-GM12878_enriched_sites BCLAF1-GM12878_enriched_sites CTCF-GM12878_enriched_sites DNase1-GM12878_enriched_sites EBF1-GM12878_enriched_sites Egr1-GM12878_enriched_sites ELF1-GM12878_enriched_sites ETS1-GM12878_enriched_sites Gabp-GM12878_enriched_sites H2AZ-GM12878_enriched_sites H3K27ac-GM12878_enriched_sites H3K27me3-GM12878_enriched_sites H3K36me3-GM12878_enriched_sites H3K4me1-GM12878_enriched_sites H3K4me2-GM12878_enriched_sites H3K4me3-GM12878_enriched_sites H3K79me2-GM12878_enriched_sites H3K9ac-GM12878_enriched_sites IRF4-GM12878_enriched_sites Jund-GM12878_enriched_sites MEF2A-GM12878_enriched_sites MEF2C-GM12878_enriched_sites Nrsf-GM12878_enriched_sites p300-GM12878_enriched_sites Pax5-GM12878_enriched_sites Pbx3-GM12878_enriched_sites PolII-GM12878_enriched_sites PolIII-GM12878_enriched_sites POU2F2-GM12878_enriched_sites PU1-GM12878_enriched_sites Rad21-GM12878_enriched_sites RXRA-GM12878_enriched_sites SIX5-GM12878_enriched_sites SP1-GM12878_enriched_sites Srf-GM12878_enriched_sites TAF1-GM12878_enriched_sites Tcf12-GM12878_enriched_sites Tr4-GM12878_enriched_sites USF1-GM12878_enriched_sites Yy1-GM12878_enriched_sites ZBTB33-GM12878_enriched_sites ZEB1-GM12878_enriched_sites. The annotated VCF files generated by SnpEff were processed using custom scripts to compute the fraction of SNPs in circQTL and background eQTL data sets, odds ratio (OR), 95% CI and p-value of the enrichment for each regulatory element. The Monte Carlo P-value for the enrichment or underrepresentation of each regulatory mark was computed as described in the functional enrichment annotation section.

### Enrichment of circQTL SNPs among GWAS loci

The NHGRI-EBI catalog of published genome wide association studies (GWAS) was downloaded from EBI (http://www.ebi.ac.uk/gwas; gwas_catalog_v1.0.1-associations_e91_r2018-01-01.tsv file). The genomic loci were separately created for circQTL and randomly generated background eQTL SNPs based on their LD patterns. To estimate LD patterns within a genomic region, PLINK [55] version 1.9 was run with the options --r2 --ld-window-kb 1000 --ld-window-r2 0.8 and SNPs with r2 > 0.8 with an index SNP were grouped together into a genomic locus. The genomic loci created for circQTL and background eQTL SNPs were sorted by chromosome and start position and converted into BED files. Similarly, columns DISEASE/TRAIT, REGION, CHR_ID, CHR_POS, MAPPED_TRAIT and MAPPED_TRAIT_URI were extracted from GWAS catalog file and processed into a BED file sorted by CHR_ID and CHR_POS. Finally, bedtools intersect [56] was used to screen overlaps of GWAS SNPs with genomic loci BED files. An associated genomic locus for a trait as defined by the EFO[40] tag of MAPPED_TRAIT_URI field in the GWAS catalog is identified if a GWAS risk SNP falls within the locus. Finally, the enrichment of circQTL SNPs among associated loci is evaluated by a one-tailed Fisher’s exact test with the following 2 × 2 table: columns; circQTL SNPs and control eQTL SNPs, rows; SNPs within and not within the disease-associated loci. A total of 306 traits were evaluated for the enrichment and the P-values obtained from Fisher’s exact test were corrected for the multiple testing based on Bonferroni correction.

## Supporting information

## ACKNOWLEDGMENTS

This work was funded by grants from Basic Medical Research Program (BMRP) from Qatar Foundation to WCM-Q.

## DECLARATIONS

### Competing interests

The author(s) declare(s) that they have no competing interests

### Data Availability

All data generated or analysed during this study are included in this published article (and its supplementary information files).

