## Supplementary figures and images for "Identification of human genetic variants controlling circular RNA expression"

### Supplementary file 1

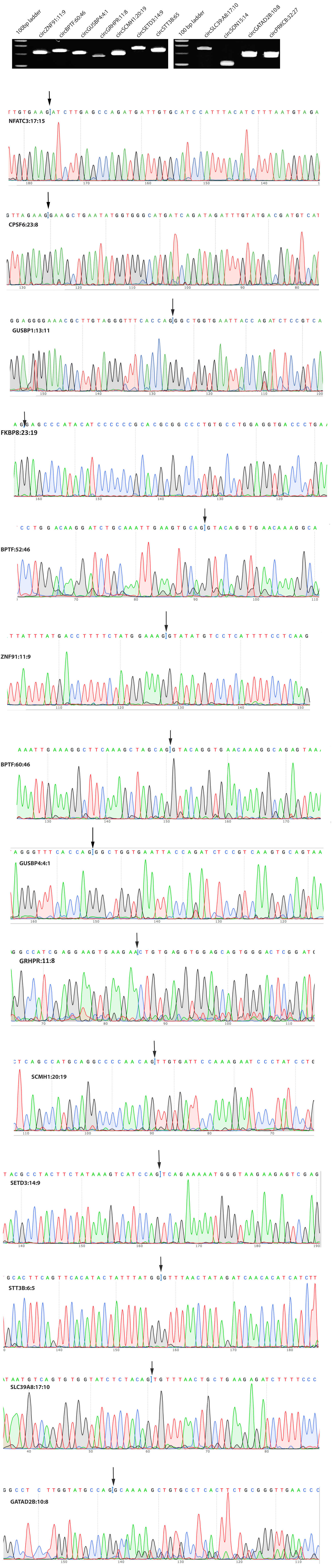
