## Supplementary material for "Identification of human genetic variants controlling circular RNA expression"

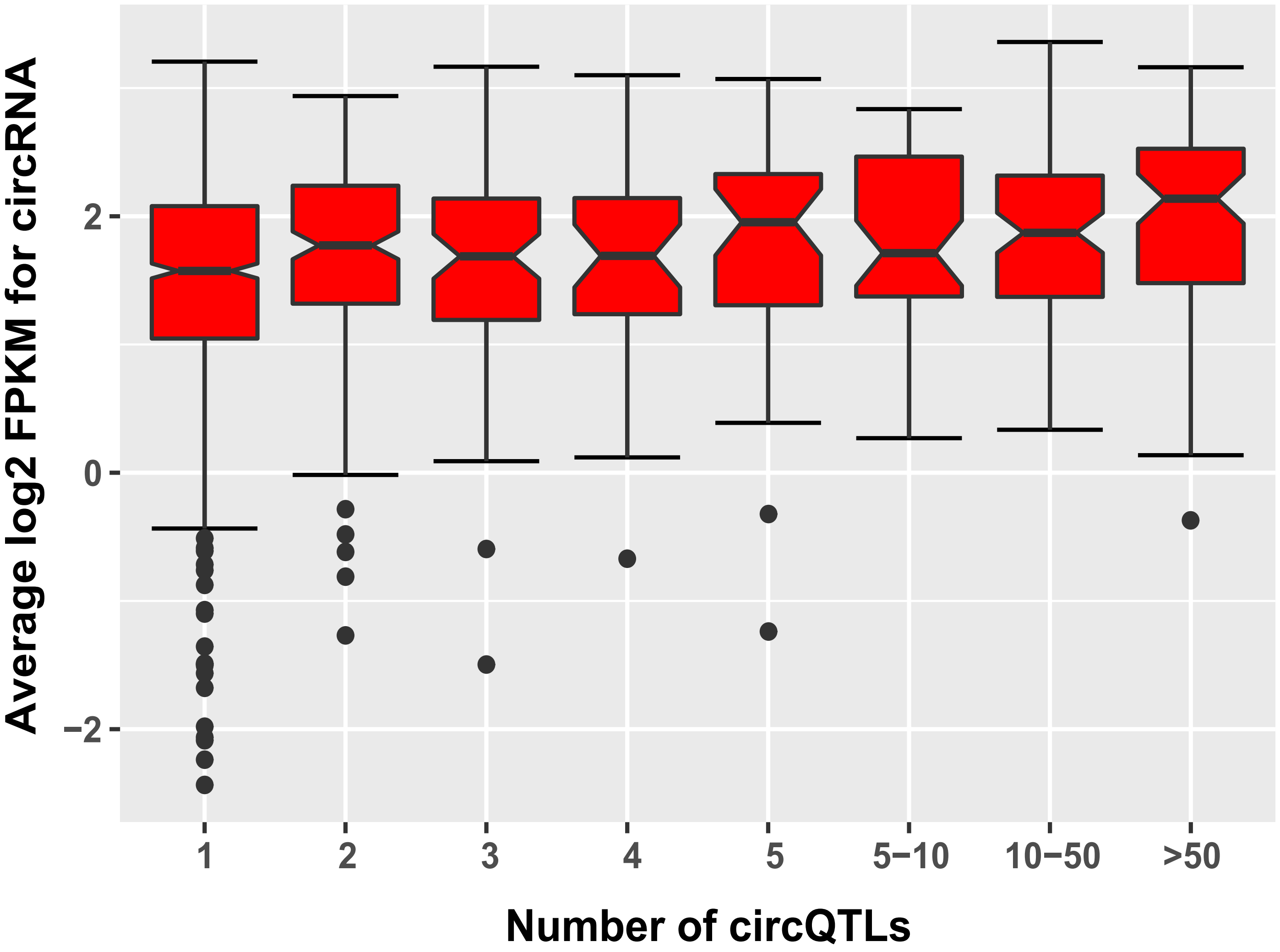


**Supplementary Figure 2: Relative Expression of circRNAs vs number of circQTLs detected.**
**Boxplots of Average circRNA expression from 358 EUR samples.**
**The data is categorized as per the number of circQTLs identified for each gene.**
