## Supplementary material for "Identification of human genetic variants controlling circular RNA expression"

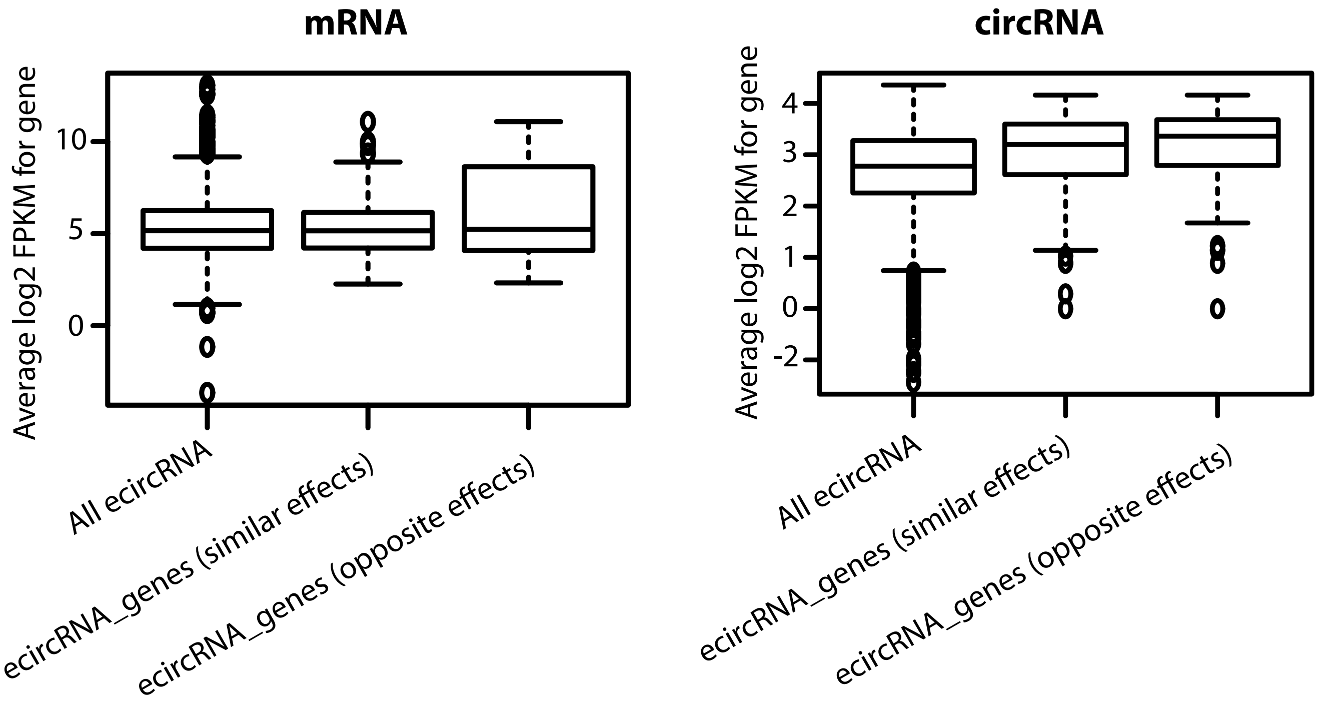


p-values from Welch Two Sample t-test for mean expression of ecircRNA_genes (opposite effects) greater than the compared category

| | **mRNA** | | **circRNA** | |
|---|---|---|---|---|
| | ecircRNA_genes (similar effects) | All ecircRNA | ecircRNA_genes (similar effects) | All ecircRNA |
| ecircRNA_genes (opposite effects) | 0.09645 | 0.05654 | 0.01095 | 3.53E-16 |

**Supplementary Figure 3: Comparison of mRNA and circRNA expression for genes containing shared QTLs.** The gene expression is evaluated to test whether genes containing shared QTLs associated with opposite effects on mRNA and circRNA expression are expressed at similar scales compared to other ecircRNA genes. The genes containing shared QTLs were divided into two categories - ecircRNA_genes (similar effects), represents genes where shared QTLs modulate circRNA and mRNA expression in a similar fashion. ecircRNA_genes (opposite effects), genes where shared QTLS have opposite effects on the circular and linear expression of their host gene.
